# Rapid 3D enhanced resolution microscopy reveals the dynamics of cortical dendritic spinules

**DOI:** 10.1101/613992

**Authors:** CR Zaccard, K Myczek, MD Martin-de-Saavedra, P Penzes

## Abstract

Dendritic spinules are thin, membranous protrusions formed by neuronal dendritic spines that are not adequately resolved by diffraction-limited light microscopy. Hence, our understanding of spinules is inferred predominantly from fixed-tissue electron microscopy (EM). Super-resolution modalities have enabled live-cell nanoscopic imaging, but their utility for fast, time-lapse, volumetric imaging has been restricted. Herein, we utilized rapid structured illumination microscopy (SIM) and ‘enhanced resolution’ confocal microscopy to study spatiotemporal spinule dynamics in live cultured cortical pyramidal neurons. Spinules on mushroom spines typically recurred at the same topographical locations and most were short-lived, originating near simple post-synaptic densities (PSDs), while a subset was long-lived and elongated, emerging from complex PSDs. Ca^2+^ puncta within spinules synchronized with spine head transients and Ca^2+^ depletion drastically decreased spinule number. Moreover, we uncovered evidence of differential Ca^2+^-mediated regulation of short-lived and long-lived spinules. Thus, we identified unique spinule classes divergent in lifespan, dynamics, morphology, relationship to the PSD, and regulation. These data suggest distinct synaptic functions of spinule classes, informing future studies, while demonstrating a new application for enhanced resolution microscopy.

## Introduction

Dendritic spinules, first detected in hippocampal neurons using EM (Westrum and Blackstad, 1962, Tarrant and Routtenberg, 1977), are thinner than filopodia, <1 µm in length, often detected on mushroom spines, and inducible by synaptic activity (Petralia et al., 2015). These nanoscale structures are not well-resolved by diffraction-limited light microscopy, hence our understanding of spinule structure and function has been largely inferred from fixed tissue EM (Tao-Cheng et al., 2009, Spacek and Harris, 2004) and a few limited-resolution confocal and two-photon imaging studies (Richards et al., 2005, Ueda and Hayashi, 2013). Prior work has established that spinules can project into pre-synaptic boutons or glial cell membranes and their proposed functions include post-synaptic membrane remodeling and retrograde signaling (Spacek and Harris, 2004, Petralia et al., 2015).

Numerous EM studies have reported their link to the PSD, including some showing spinule origination from PSD edges or perforations (Tao-Cheng et al., 2009, Spacek and Harris, 2004, Petralia et al., 2015), and others describing origination from nonperforated PSD edges or spine necks (Sorra et al., 1998). A study utilizing fast, multi-channel, high-resolution imaging in live neurons could provide needed clarification on the relationship between spinules, which can be highly dynamic (Tao-Cheng et al., 2009), and PSDs, which are also continually remodeling in response to activity (Okabe et al., 1999, Bourne and Harris, 2008). Moreover, Ca^2+^ is a key signaling ion in a multitude of synaptic functions, including synaptic transmission, which is known to induce spinule formation (Petralia et al., 2015, Tao-Cheng et al., 2009, Ueda and Hayashi, 2013). Ca^2+^ regulates spinule-dependent synaptic plasticity in fish horizontal cells (Country and Jonz, 2017), but the role of Ca^2+^ in mammalian spinule formation remains unclear, necessitating live, high-speed imaging studies.

Recently developed super-resolution imaging modalities have enabled live imaging of nanoscale structures (Cox, 2015, Godin et al., 2014), but their application for large volumetric datasets has been restricted by slow acquisition speed, photo-bleaching, and photo-toxicity. While SIM is well-suited for live three-dimensional (3D) imaging, a mechanically translated grating pattern has restricted acquisition speed, leading to missed events and motion artifacts (Fiolka et al., 2012). Here we utilized a custom-built system that employs a liquid crystal spatial light modulator to efficiently alter the grating pattern and increase speed (Kner et al., 2009, Li et al., 2015) to quantify spinule lifespan and dynamics. We additionally used enhanced resolution confocal microscopy to investigate spinule dynamics in relation to the PSD, as well as Ca^2+^ as a candidate regulator of mammalian spinule formation during basal activity.

## Results

We first utilized a SIM system with improved optical sectioning speed compared to standard SIM to image live dissociated cortical mouse pyramidal neurons. We balanced acquisition speed against photobleaching and phototoxicity, to resolve dynamic, thin membrane protrusions that were finer than filopodia and originated from dendritic spines, i.e. spinules, at 15–20 sec intervals. Similar to a previous report (Spacek and Harris, 2004), IMARIS 3D reconstructions revealed spinules on mushroom, branched, and more rarely, on thin spines (Figure 1A). Mushroom spines displayed a substantially higher mean number of spinules over the 1000 sec imaging duration, compared to spines transitioning between classes, thin, or filopodia-shaped spines (Figure 1B). We next tracked mushroom spine volume over time (Figure 1C) and found a strong positive correlation between spinule number and mean spine head volume (Figure 1D). Spinules were often observed in non-contiguous frames at the same topographical spine head locations, and repeatedly extending and retracting along axons (Figure 1E; Video 1). Quantification of normalized spinule origination in relation to kymograph outlines of spine heads (Figure 1F) showed that the majority (76%) of spinules recurred within the 1000 sec imaging frame, and the mean number of recurring spinules per spine was substantially higher than singly occurring spinules (Figure 1G). Importantly, most spinules (70%) existed for <60 sec, while the minority (30%) existed for ≥60 sec, rarely exceeding the imaging duration (Figure 1H and Figure S1A). Short-lived spinules were accordingly more frequent per spine (Figure 1I). A representative spine displayed multiple short-lived spinules, which were often tapered or vermiform, and fewer longer-lived spinules (Figure 1J; Figure S1B-C; Video 2). Long-lived spinules also displayed diverse morphologies, including pinch-waisted and rarely filopodia-like, as well as occasional spine-neck origination and evidence of presumptive vesicular trafficking within thick spinules (Figure 1K, Figure S1D).

**Figure 1.**
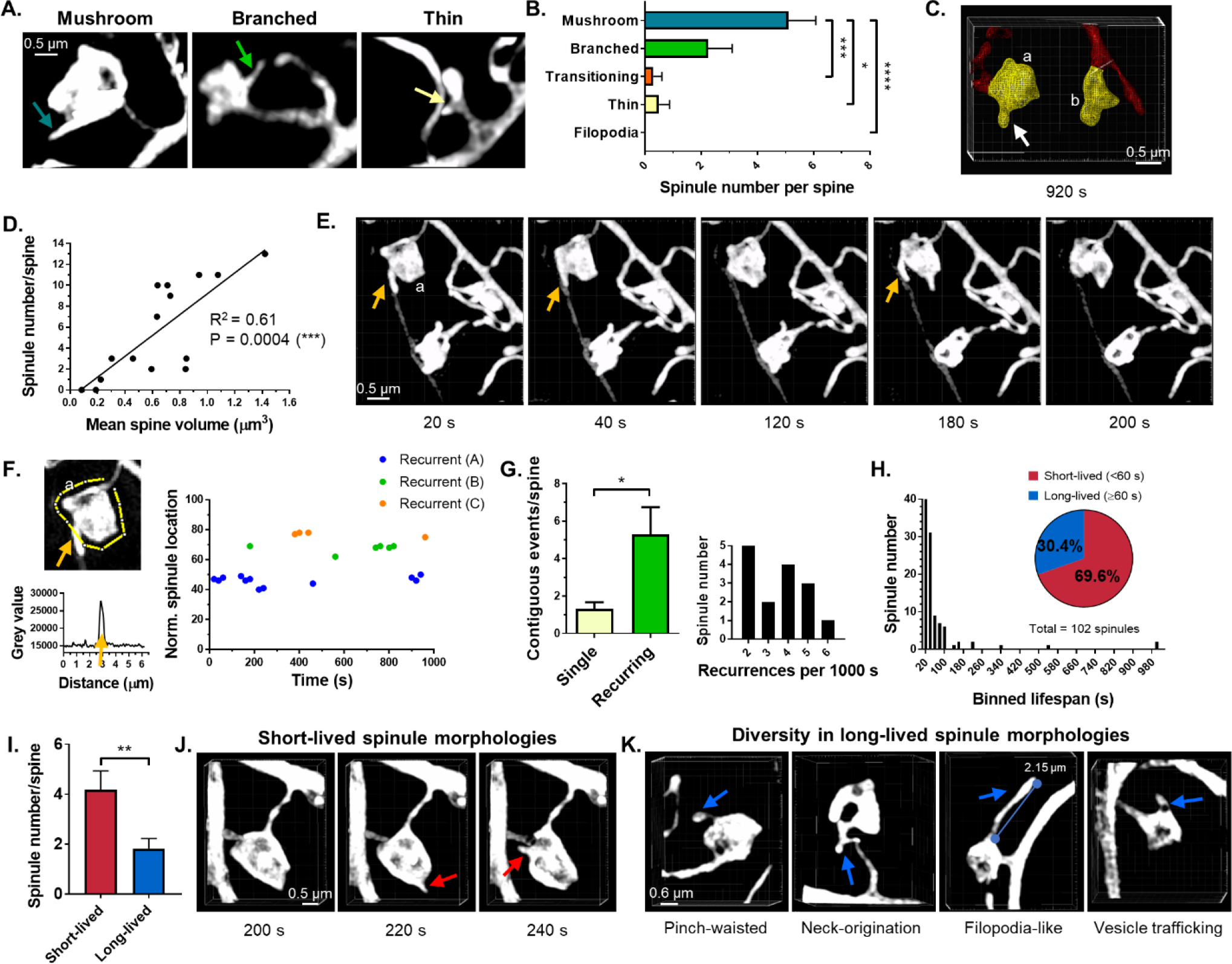
Dendritic spinule recurrence and divergent lifespans. (**A**) Representative images of spine morphological subtypes displaying spinules (arrows). (**B**) Spinule frequency on spine subtypes (n=54 spines; mushroom *vs.* transitioning P=0.0009; mushroom *vs.* thin P=0.0153; mushroom *vs.* filopodia P<0.0001). (**C**) Wire-frame IMARIS 3D reconstructions of a representative large-volume (a; mean=0.64 µm^3^; 10 spinules) and small-volume mushroom spine (b; mean=0.22 µm^3^; one spinule). (**D**) Positive correlation between spinule number and mushroom spine volume (n=15 spines). (**E**) Montage highlighting spinule extension and retraction (arrows) along axon, and recurrence at one topographical location. (**F**) Normalized temporal location mapping showing recurrent spinules from spine (a). (**G**) Frequency of single and recurring spinules (n=66 spinules; P=0.0004). (**H**) Frequency distribution of binned spinule lifespans and pie chart of spinule subgroups (n=102 spinules). (**I**) Short-lived compared to long-lived spinule number per spine (n=17 spines). (**J**) Characteristic short-lived spinule morphologies (red arrows). (**K**) Diversity of long-lived spinule morphologies. Data are represented as mean ± SEM. *P<0.05, **P<0.01, ***P<0.001, ****P<0.0001.

We next assessed spinule dimensions over time using rapid SIM. The current consensus is that these structures are typically < 1 µm in length and finer than filopodia, while reports on their diameter and shape have varied (Petralia et al., 2015). We found that the mean length of all spinules over their lifespans was 0.49 µm, while the mean width and volume was 210 nm and 38 nm^3^, respectively (Figure S2A-D). Furthermore, mean spinule length positively correlated with lifespan (Figure 2A), but not width or volume (DNS). Long-lived spinules were significantly greater in mean and maximum length (Figure 2B), with no difference in mean width between groups (Figure S2E). We tracked spinule volumes using IMARIS to generate temporal spinule volume plots (Figure 2C) and illustrate the differential size and stability of short-versus long-lived spinules (Figure 2D-E; Video 3). Overall, mean and maximum volume was greater in long-lived than short-lived spinules (Figure 2F). Strikingly, the mean spinule length change was negatively correlated with spinule lifespan (Figure 2G). While the cumulative length and volume change was greatest in long-lived spinules, the change in length (nm/sec) and volume (nm^3^/sec) was dramatically higher in short-lived spinules compared to long-lived spinules (Figure 2H-I; Figure S2F-G). These data suggest the existence of distinct spinule classes with divergent lifespans, morphologies, and dynamics.

**Figure 2.**
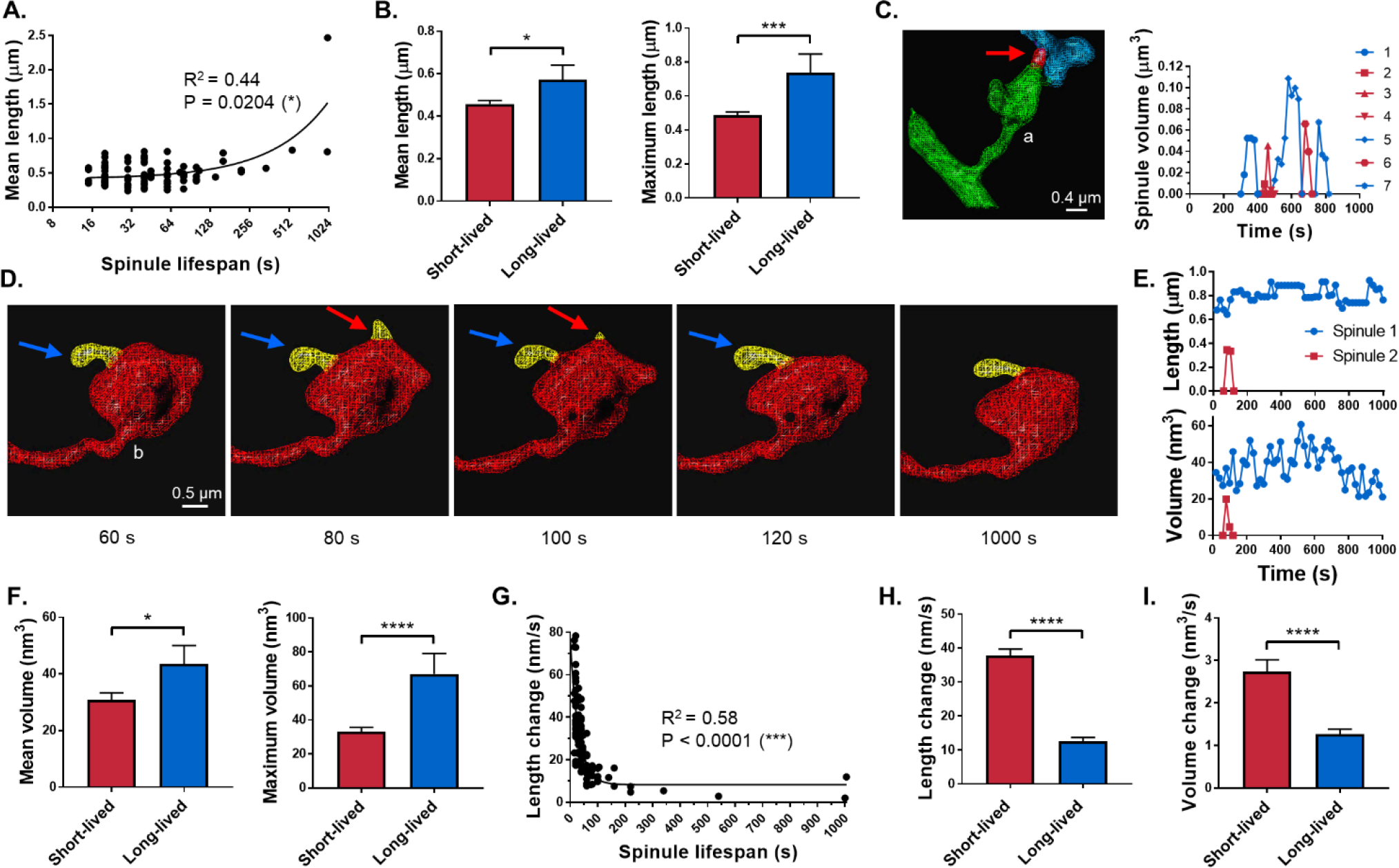
Differential short- and long-lived spinule dimensions and dynamics. (**A**) Correlation between mean spinule length and lifespan (n=102 spinules). (**B**) Mean and maximum length of short-*vs.* long-lived spinules (mean P=0.0436; max. P=0.0002). (**C**) Wire-frame 3D model showing single-frame spinule volume, and corresponding plot of short-(red) and long-lived spinule (blue) volumes over time. (**D**) Montage of a spine displaying a long-lived, pinch-waisted spinule (blue arrows) and formation and dissolution of a tapered, short-lived spinule (red arrows). (**E**) Temporal maps of spinule lengths and volumes corresponding to spine b, showing long-lived spinule stability. (**F**) Mean and maximum volume in short-*vs.* long-lived spinules (n=84 spinules; mean P=0.0159; max. P<0.0001). (**G**) Negative correlation between mean spinule length change and lifespan (n=102 spinules). (**H**) Mean change in length of spinules grouped by lifespan (n=102 spinules; P<0.0001). (**I**) Mean volume change in short-*vs.* long-lived spinules (n=84 spinules; P<0.0001). Data are shown as mean ± SEM. *P<0.05, **P<0.01, ***P<0.001, ****P<0.0001.

Some studies have reported spinules originating from PSD edges or perforations (Tao-Cheng et al., 2009, Spacek and Harris, 2004, Petralia et al., 2015), versus nonperforated PSD edges or spine necks (Sorra et al., 1998). Considering that PSDs too are continually remodeling in response to activity (Okabe et al., 1999, Bourne and Harris, 2008), fast, multi-channel, high-resolution imaging in live neurons is needed to clarify the link between spinule origin and the PSD. Hence, we utilized fluorescently-tagged ‘intrabodies’ to label endogenous PSD95 (Gross et al., 2013) and resonance scanning enhanced resolution confocal microscopy (Figure 3A), to mitigate intrabody photo-bleaching observed with SIM, which requires a strong fluorescence signal for time-lapse imaging. The majority (91%) of mushroom spines detected by this alternate method were spinule-positive. Rare, elongated, long-lived spinules and frequent, short-lived spinules were again detected over 600 sec, and the shorter frame interval enabled the detection of an extremely short-lived subgroup (<20 sec) (Figure S3A-B). In accordance with rapid SIM, a significant positive correlation was found between mean spinule length and lifespan, and both the mean and maximum spinule length was substantially higher in long-as opposed to short-lived spinules (Figure S3C-D). Short-lived spinules typically originated near to the PSD edge, while long-lived spinules originated substantially further from the edge of more complex PSDs, at a mean distance of 0.28 and 0.43 µm, respectively (Figure 3B-D; Videos 4 and 5). Surprisingly, PSD volume and dynamics, including the change in PSD volume per sec and the net +/-volume change, did not correlate with spinule number per spine (Figure 3E-F; Figure S3E-F). Complementing prior EM findings, long-lived spinules occasionally contained mobile PSD fragments that could traverse their length (Figure 3G; Video 6). Strikingly, 78% of spines displaying long-lived spinules also contained complex, partitioned PSDs compared to 23% of long-lived spinule-negative spines (Figure 3H), and 67% of long-lived spinule-positive spines displayed fragmented PSDs versus 28% of long-lived spinule-negative spines (Figure 3I). Mean and maximum PSD fragment number per frame was significantly greater in long-lived spinule-positive compared to long-lived spinule-negative spines (Figure 3J). These data suggest a link between complex PSD remodeling and spinule stability, while simple PSDs are associated with dynamic, transient spinules.

**Figure 3.**
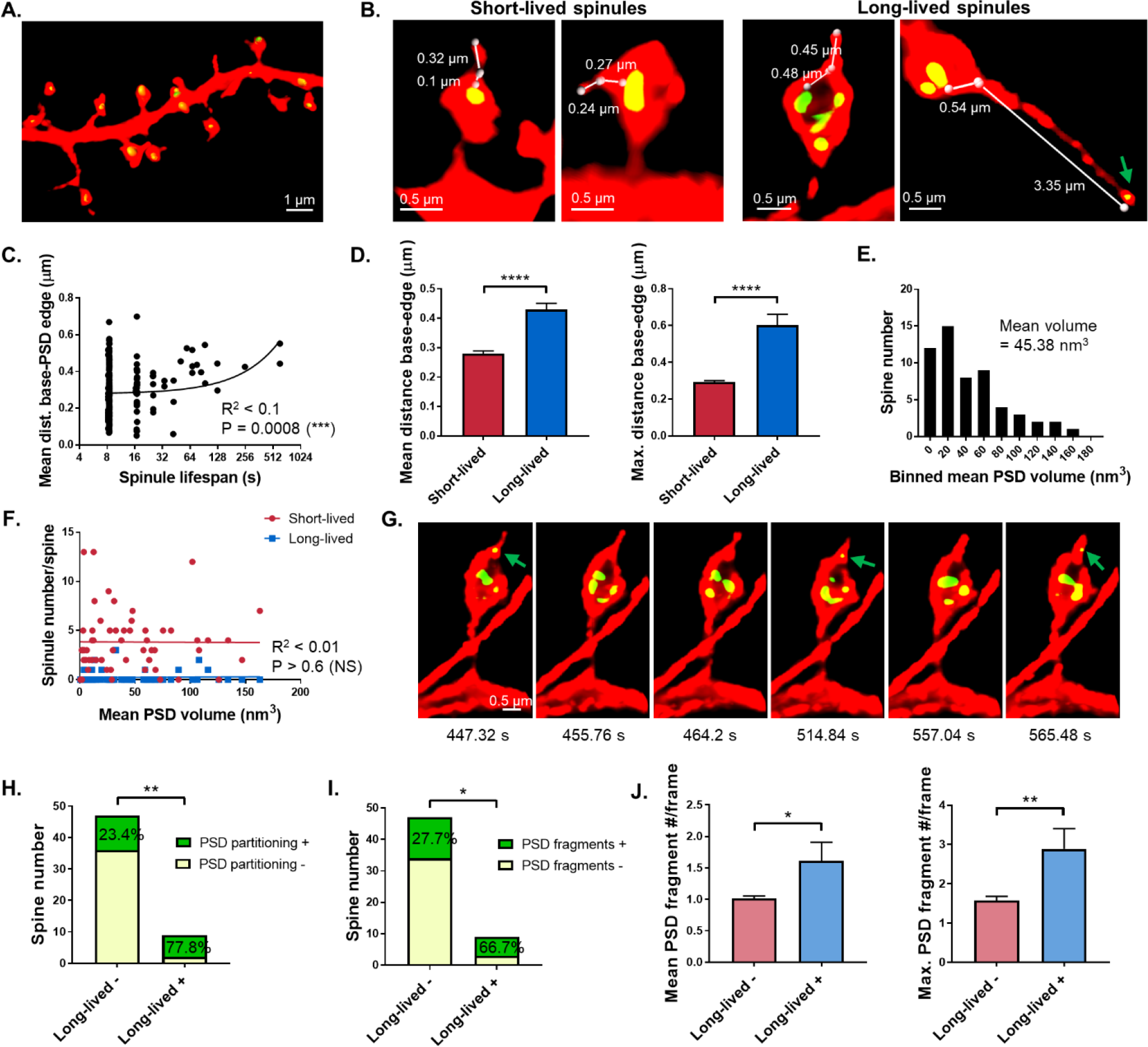
Relationship between spinule subgroups and PSD complexity. (**A**) Endogenous PSD95 labeling with GFP-tagged intrabodies followed by enhanced resolution confocal imaging and IMARIS analysis. (**B**) Representative spines with short-lived spinules originating near the PSD edge compared to long-lived spinules originating farther from the PSD edge. PSD fragment was visualized in a retracting filopodia-like spinule tip. (**C**) Correlation between mean distance from spinule base to PSD edge and spinule lifespan (n=228 spinules). (**D**) Mean and maximum distance from base to PSD edge in short-lived *versus* long-lived spinules (mean P<0.0001; max. P<0.0001). (**E**) Binned frequency distribution of mean PSD volume per spine over time (n=56 mushroom spines). (**F**) No correlation between spinule number and mean PSD volume. (**G**) Montage of small PSD fragment trafficking into a long-lived spinule. (**H**) Proportion of long-lived spinule-negative and -positive spines displaying substantial PSD partitioning (n=56 spines; P=0.0032). (**I**) Proportion of long-lived spinule-negative and -positive spines displaying simple *versus* fragmented PSDs (n=56 spines; P=0.0491. (**J**) Mean and maximum PSD fragment number per frame in long-lived spinule-negative and -positive spines (n=56; mean P=0.0388; max. P=0.0045). Data are represented as mean ± SEM. *P<0.05, **P<0.01, ***P<0.001, ****P<0.0001.

Ca^2+^ is a key signaling ion in synaptic transmission, which induces spinule formation (Petralia et al., 2015, Tao-Cheng et al., 2009, Ueda and Hayashi, 2013), and regulates spinule-dependent synaptic plasticity in the retina of teleosts (Country and Jonz, 2017). To investigate Ca^2+^ as a candidate regulator of mammalian spinules during basal activity, we again utilized fast (2.2 sec/frame) enhanced resolution confocal imaging of neurons expressing the Ca^2+^ sensor GCaMP6s (Chen et al., 2013). We again observed a preponderance of short-lived and infrequent long-lived spinules in negative control spines, comparable to prior enhanced resolution confocal imaging data (Figure S4A). Interestingly, we visualized isolated Ca^2+^ puncta appearing in long-lived spinules repeatedly during Ca^2+^ transients in control spines (Figure S4A; Video 7). Plots of GFP mean fluorescence intensity (MFI) in the dendritic shaft, spine head, and spinules of controls also revealed synchronization of Ca^2+^ transients in spinules with that of spine heads and dendritic shafts (Figure 4B; Figure S4B). To evaluate the requirement for intracellular Ca^2+^ in spinule formation, we compared vehicle-treated control neurons to those treated with the cell-permeant Ca^2+^ chelator, BAPTA-AM. The total number of short- and long-lived spinules and cumulative spinule lifespan were significantly higher in controls than in BAPTA-AM-treated spines (Figure 4C). To assess the relationship between patterns of Ca^2+^ transients and spinule dynamics, we first determined the change in Ca^2+^ fluorescence (ΔF/F_0_) in dendritic shafts and spine heads, and then generated corresponding spatio-temporal maps to track spinule number and recurrence over time. Mushroom spines with frequent, high-amplitude Ca^2+^ transients displayed many short-lived and fewer long-lived spinules (Figure 4D-E; Video 8), while spines with sporadic peaks formed only short-lived spinules (Figure S4C-D). In contrast, spines from BAPTA-AM-treated neurons displayed greatly dampened Ca^2+^ transients and substantially fewer spinules (Figure 4F-G; Video 9). Quantitative assessment of Ca^2+^ peaks in spines, i.e. 50% increase in ΔF/F_0_ above baseline, revealed a positive correlation between total spinule number and peak amplitude (Figure S4E), but no correlation with peak frequency or duration. Interestingly, the number of short-lived, but not long-lived, spinules per spine positively correlated with Ca^2+^ peak mean and maxima (Figure 4H). Conversely, long-lived spinule number positively correlated, albeit less strongly, with Ca^2+^ peak frequency and duration. These findings suggest distinct mechanisms of spinule regulation by local Ca^2+^ transients, informing future functional studies of spinule subgroups.

**Figure 4.**
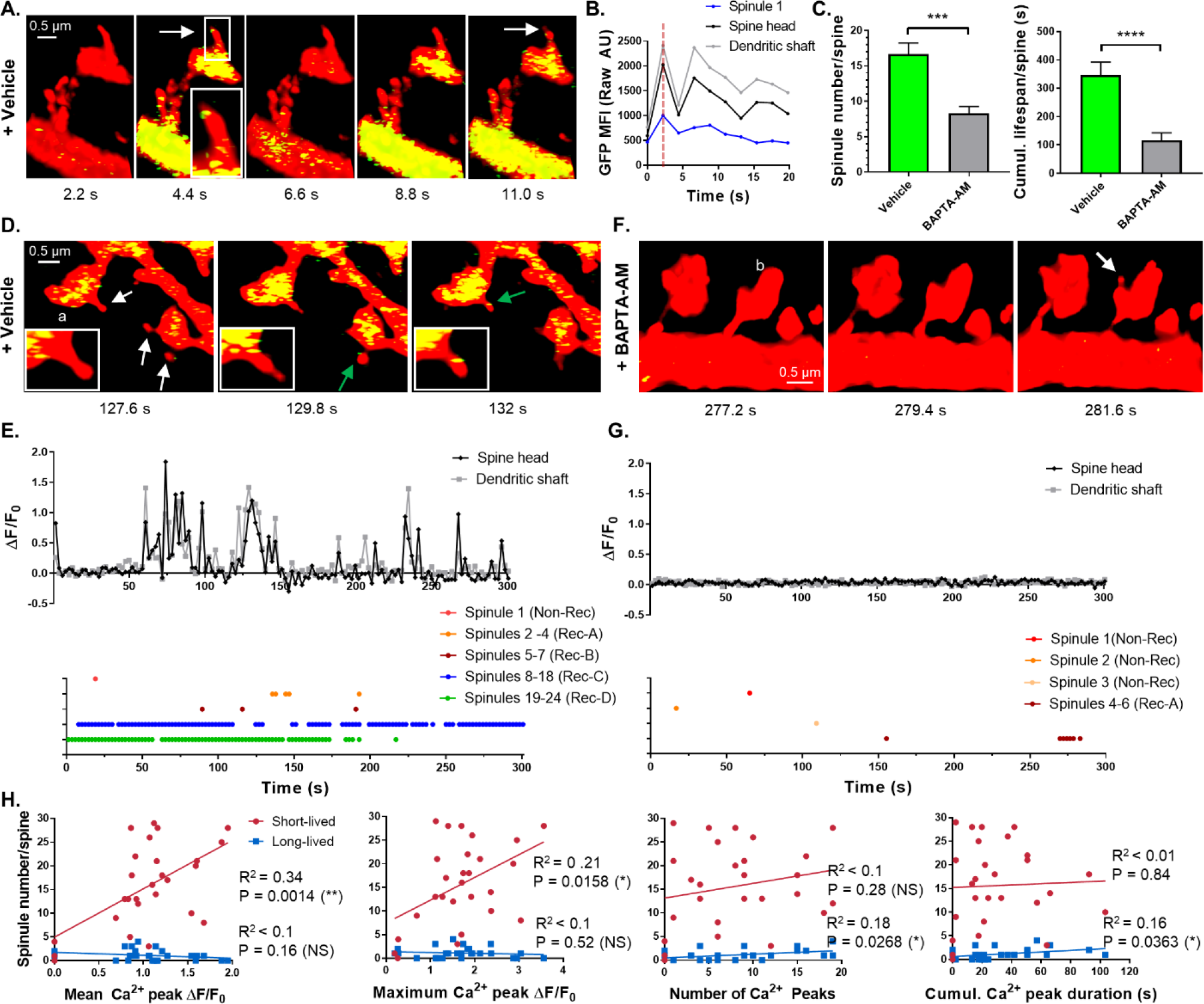
Differential regulation of spinule subgroups by Ca^2+^. (**A**) Vehicle-treated control spine montage showing Ca^2+^ puncta localized to a long-lived spinule tip during spine transients. (**B**) Synchrony of Ca^2+^ activity in spinule, spine head, and dendritic shaft. (**C**) Comparison of spinule number and cumulative lifespan in spines from vehicle-*versus* BAPTA-AM-treated cells (vehicle, n=28, P=0.0001; BAPTA-AM, n=28 spines, P<0.0001). (**D**) Ca^2+^ puncta entering a long-lived spinule synchronized with Ca^2+^ peaks in control spine a. (**E**) ΔF/F_0_ ratio in spine head a and dendritic shaft over time and corresponding recurring spinule map. (**F**) BAPTA-AM-treated spines displaying dampened Ca^2+^ transients and few spinules. (**G**) ΔF/F_0_ ratio in spine head b and dendritic shaft and corresponding temporal recurrent spinule map. (**H**) Correlations between short- and long-lived spinule number per spine and Ca^2+^ peak amplitude, frequency, or duration (n=27 spines). Data are represented as mean ± SEM. *P<0.05, **P<0.01, ***P<0.001, ****P<0.0001.

## Discussion

Herein, we demonstrate the utility of two complementary enhanced resolution live imaging techniques for studying dynamic nanoscale post-synaptic membrane projections in volumetric samples. While rapid SIM was ideal for distinguishing fine structural details at moderate acquisition speeds, enhanced resolution was the optimal choice for faster, multi-channel imaging, particularly when the fluorescence signal is weak, while still providing the resolving power necessary for spinule visualization. We observed by rapid SIM diverse spinule morphologies, which mirrored those seen in slice cultures by EM (Tao-Cheng et al., 2009). Our findings also provide the first evidence of spinule recurrence, suggesting localized molecular machinery at spinule origination sites, and of distinctive classes of short-lived, dynamic spinules and fewer long-lived, stable spinules.

While spinules have been proposed to participate in post-synaptic membrane remodeling and PSD assembly (Spacek and Harris, 2004), their dynamic relationship with the PSD has not been investigated in living neurons. Our findings that short-lived spinules typically originate near to simple PSDs, while long-lived spinules originate farther from complex PSDs, often extending from perforations, imply a relationship predominantly between long-lived spinules and PSD remodeling. The distinctive morphologies and dynamics of spinule classes together suggest divergent modes of regulation and functions. Long-term potentiation is known to induce spinule formation (Ueda and Hayashi, 2013, Petralia et al., 2015), but the role of Ca^2+^ in mammalian spinule formation had not been investigated. Our results strongly indicate that Ca^2+^ is involved in their regulation. We also observed Ca^2+^ puncta within spinules, isolated from, but in synchrony with spine heads, raising the question of whether Ca^2+^ enters spinules from the spine head or via channels on the spinule surface. Interestingly, the nuanced regulation of short-lived and long-lived spinules by Ca^2+^ transients in spines also suggests distinct synaptic functions. In summary, we demonstrate a new application of enhanced resolution microscopy to study dynamic sub-spine membrane projections, informing ongoing mechanistic and functional studies.

## Acknowledgments

SIM was performed at the Advanced Imaging Center of the Janelia Research Campus, funded by the Howard Hughes Medical Institute and Gordon & Betty Moore Foundation. Confocal microscopy was performed at Northwestern’s Nikon Imaging Center with the support of Drs. Constadina Arvanitis and David Kirchenbuechler. This work was supported by NIH grants, R01MH107182 and R01MH071316.

## Author contributions

PP conceived and supervised the project, and CRZ designed and conducted experiments with the assistance of KM and MDM.

---

**Figure S1.**
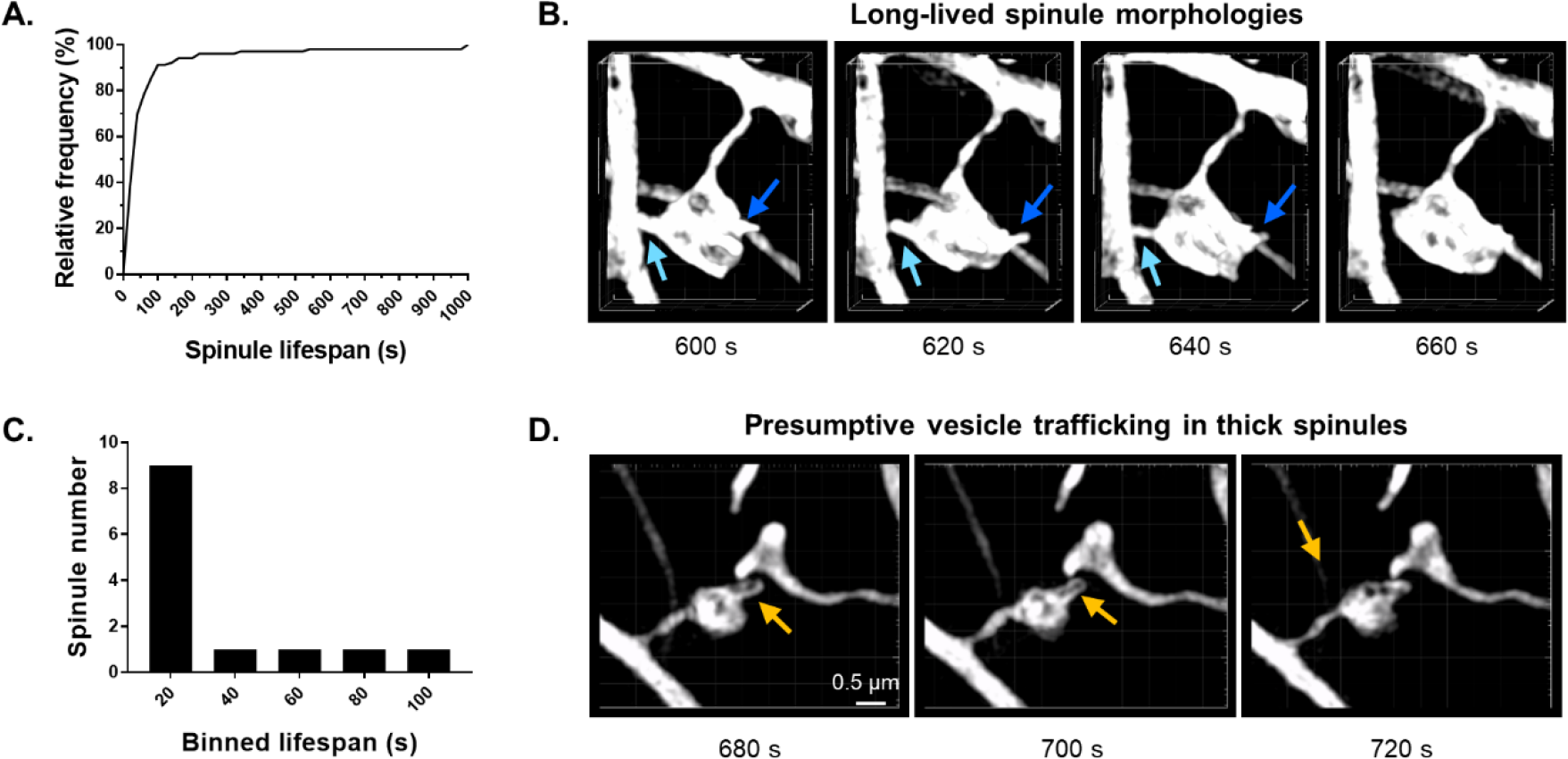
Spinule recurrence and diversity in long-lived spinule morphologies. (**A**) Relative frequency distribution (%) of all spinule lifespans that were detected in mushroom spines using rapid SIM (n=102 spinules). (**B**) Montage highlights a large mushroom spine, also shown in Fig. 1j and Supplementary Video 2, developing long-lived spinules (≥ 60 s) that are vermiform or tapered in shape (blue arrows) and contact adjacent dendritic shafts and axons. (**C**) Corresponding binned frequency distribution shows individual spinule lifespans in the spine shown previously over the imaging duration. (**D**) Montage of a mushroom spine that displays evidence of presumptive vesicular trafficking (arrows) between the spine head and a thick spinule, which simultaneously contacts an adjacent spine head.

**Figure S2.**
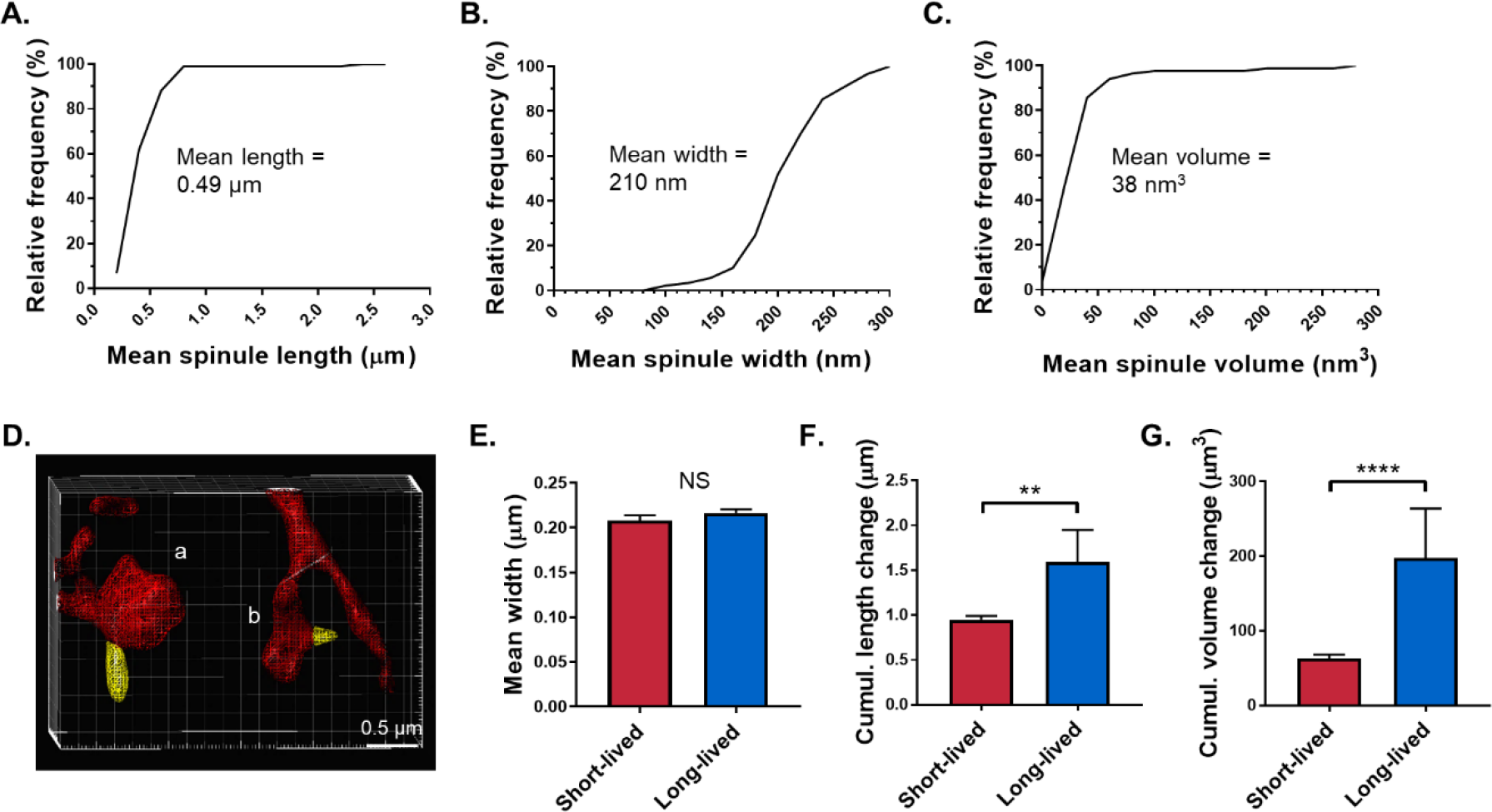
An evaluation of spinule dimensions over time. (**A**) Cumulative frequency distribution of mean spinule length (µm) over the imaging duration (n=102 spinules). (**B**) Cumulative frequency distribution of spinule width (nm) (n=89 spinules). (**C**) Cumulative frequency distribution of mean spinule volume (nm^3^) (n=84 spinules). (**D**) Wireframe model of high volume a and low volume b spines also depicted in Fig. 1C. Spinule volumes (yellow) were quantitated in each frame using IMARIS. (**E**) Spinules were grouped according to lifespan and the difference in mean width (µm) assessed between groups (n=89 spinules). (**F**) The cumulative length change, defined as the sum of the absolute value of difference in length from frame to frame, was assessed in short-compared to long-lived spinules (n=102 spinules). (**G**) The cumulative change in volume over time was compared in long-versus short-lived spinules (n = 84 spinules). Data are represented as mean ± SEM. *P<0.05, **P<0.01, ***P<0.001, ****P<0.0001.

**Figure S3.**
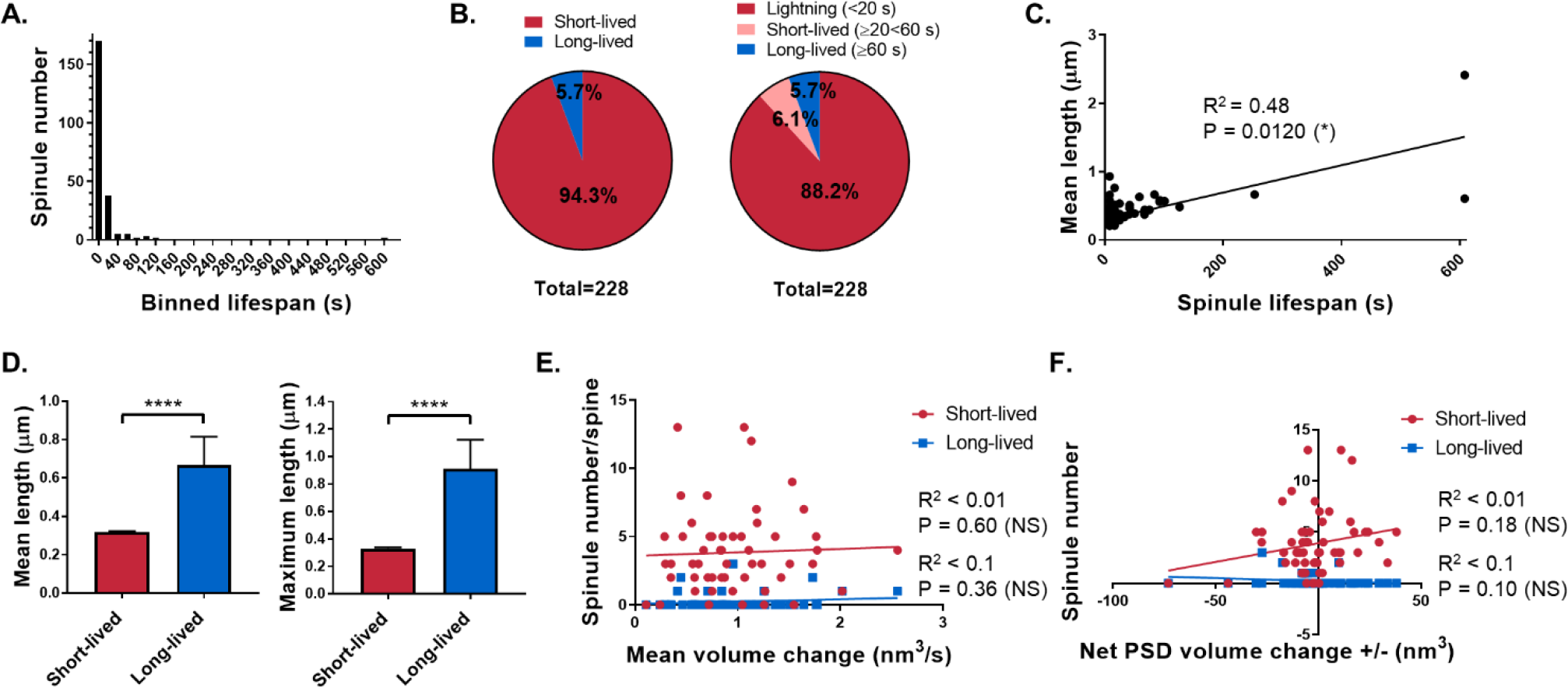
Spinule characterization relative to the PSD using enhanced resolution confocal imaging. (**A**) Binned frequency distribution of spinules on mushroom spines according to lifespan (n=228 total spinules). (**B**) Pie-charts showing the percentage of spinules classified into two groups, short- or long-lived, as before, or three groups, lightning (<20 sec), short-lived (≥20<60 sec), and long-lived (≥60 sec). (**C**) Correlation between mean spinule length and lifespan. (**D**) Mean and maximum length of short-lived versus long-lived spinules over time. (**E**) Correlation between the total number of short- and long-lived spinules per spine and mean PSD volume change. (**F**) Correlation between the number of short- and long-lived spinules per spine and the net PSD volume change (+/−) over time. Data are represented as mean ± SEM. NS≤0.05, *P<0.05, **P<0.01, ***P<0.001, ****P<0.0001.

**Figure S4.**
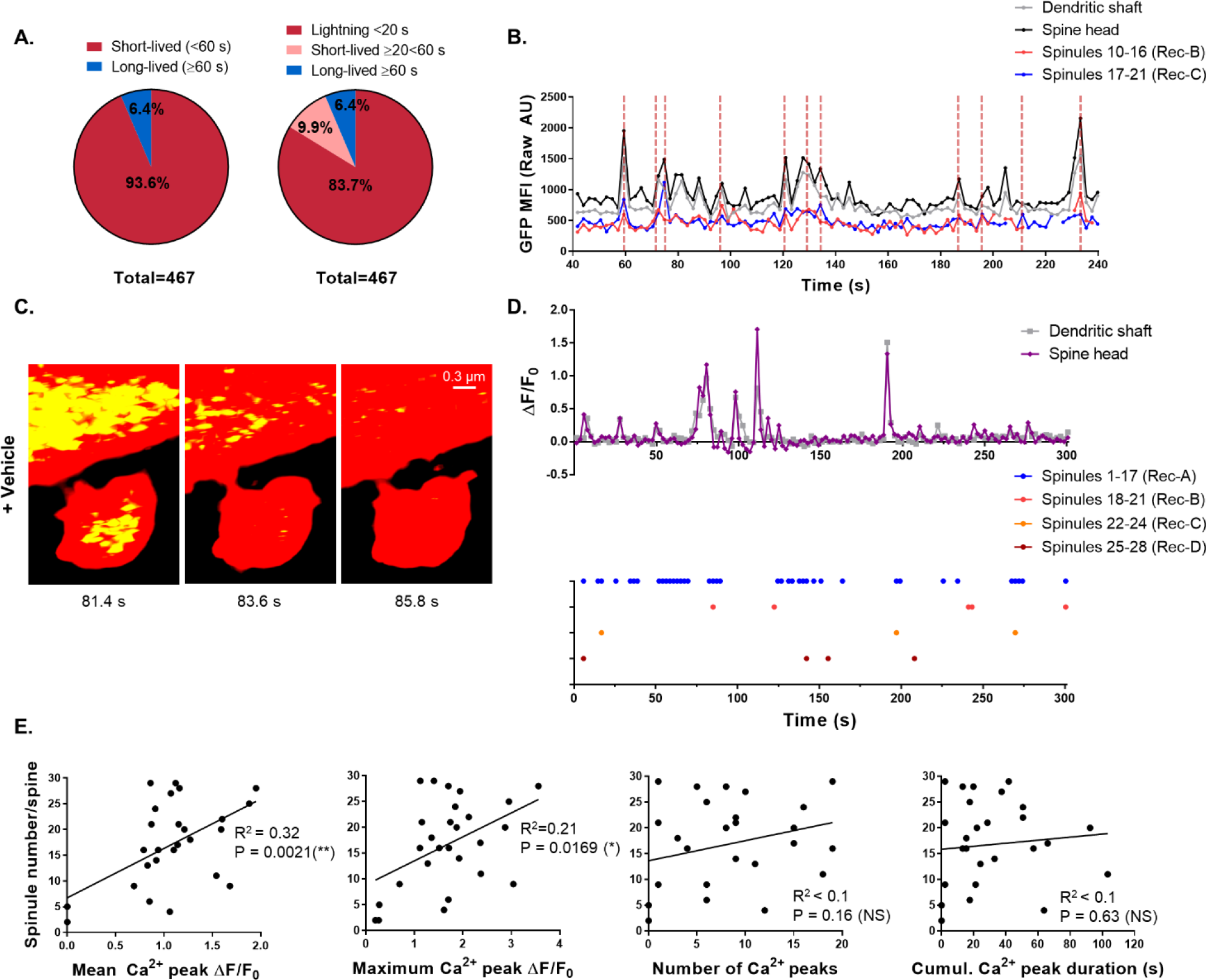
Spatiotemporal relationship between Ca^2+^ and spinules. (**A**) Pie charts showing the percentage of spinules separated into two or three groups according to lifespan detected over a 300 sec duration at 2.2 sec/frame using enhanced resolution confocal imaging (n=10 BAPTA-AM-treated cells and 27 spines; n=11 vehicle-treated cells and 27 spines). (**B**) Plot of the GCAMP6s MFI (raw AU) within ROIs of the dendritic shaft, spine head, and two recurring spinules that repeatedly form at the same topographical spine head locations over time. Dashed lines highlight Ca^2+^ peaks in spinules and their synchrony with peaks in the spine head and/or dendritic shaft. (**C**) Vehicle-treated control mushroom spine displaying an enhanced number of short-lived spinules, and a paucity of long-lived spinules. (**D**) Corresponding plot of Ca^2+^ transients expressed as ΔF/F_0_ over time in the dendritic shaft and spine head, which display low frequency, sporadic, and high-amplitude Ca^2+^ peaks. Spatio-temporal spinule map for that spine tracks four recurring spinules that reappear at the same topographical spine head locations over the imaging duration. (**E**) Correlations between the total number of spinules, including short- and long-lived spinules, per spine and Ca^2+^ peak mean amplitude, maximum amplitude, frequency, or duration.

## Supplementary Information

### Experimental procedures

#### Murine cortical dissociated neuron cultures

C57BL6 mice were bred at the Janelia Research campus in accordance with their “Use of Animals in the Advanced Imaging Center Protocol” #15-120 or at Northwestern University in accordance with animal use protocol #IS00001722. Dissociated cortical neurons were prepared from P0 mouse pups. Briefly, pups were sacrificed, and the brains were extracted and placed in cold L-15 medium (Millipore Sigma) plus Penicillin/Streptomycin (P/S) solution (Gibco). Meninges were removed with fine forceps and the cortices dissected with the aid of a dissection microscope and transferred to cold L-15 media. Media was replaced with pre-warmed 0.25% trypsin (HyClone) and the tubes were placed in 37°C H_2_0 bath for 10 min. Trypsin was replaced with RF medium (426 ml high glucose DMEM (Thermo Fisher Scientific) + 50 ml FBS (Life Technologies) + 3.5 ml P/S solution + 3.5 ml 100X L-Glutamax (Sigma) + 15 ml sterile filtered 20% D-(+)-glucose (Milipore Sigma), and cells were mechanically dissociated by gentle pipetting. Cell suspension was passed through a 40 µm nylon cell strainer and plated to 35 mm 1.5 coverslip poly-D-lysine (PDL)-coated glass-bottom imaging dishes (Mattek) that were coated a second time with PDL (0.2 mg/ml, Milipore Sigma), at a density from 1.2 × 10^6^ cells per dish. Dishes were placed in 37°C, 5% CO_2_ incubator for at one hour to allow attachment, and then media was replaced with neurobasal (NB) medium without phenol red (Gibco) + supplements, i.e. 100X Glutamax I (Gibco), B27 supplement (Life Technologies), and P/S. After four days, cultures were supplemented with 200 μM D, L-amino-phosphonovalerate (APV) (Abcam), and feeding was repeated with 100 µM APV every four days up to 16 days in culture.

#### Plasmid transfections

Chemical transfections were performed at 37°C, 5% CO_2_ in NB plus supplements media without the addition of P/S. First, 7 µl Lipofectamine 2000 (LFA2K) (Thermo Fisher Scientific) was mixed with 100 µl Opi-MEM 1X (Gibco) per imaging dish and then 50 µl of Opti-MEM containing 3.5 µg of DNA DsRedExpress2-N1 (AddGene) for rapid SIM studies and 3.5 µg mRuby-N1 plasmid (AddGene) and 3.5 µg of pCAG_PSD95.FingR-eGFP-CCR5TC (AddGene) or pGP-CMV-GCAMP6s (AddGene) for A1 confocal microscopy studies. LFA2K and plasmid DNA was mixed by pipetting up and down 30 times and incubated for 20-30 min at 37°C. The mixture was then added to neurons in antibiotic-free medium and incubated for four hours, when the media was replaced with NB plus supplements. Dishes were incubated for two days to allow for adequate expression of plasmid-encoded fluorescent proteins for filling cell structures, i.e. dendritic spines and spinules, and labeling of endogenous PSD95 or Ca^2+^ transients.

#### BAPTA-AM or vehicle treatment

Nineteen-day old mouse cortical neurons were transfected with mRuby and GCAMP6s plasmids. After two days to allow for sufficient plasmid expression, NB media was replaced with pre-warmed 1X ACSF (Smith et al., 2014) and each dish let rest for 30 min. Media was replaced with pre-warmed 1X ACSF plus 20 µm BAPTA-AM diluted in DMSO (Sigma-Aldrich) or the equivalent volume of DMSO alone (0.5 µl/ml) for the treatment and negative control group, respectively. After 30 min of treatment at 37°C in 5% CO_2_, live imaging was conducted on a single stage position per dish.

#### Microscopy and image processing

All imaging was conducted on live neurons at 21 days of age in an environmental imaging chamber at 37°C, 5% CO_2_. Image fields were selected on secondary branches of apical dendrites on excitatory pyramidal neurons that showed no signs of poor health, e.g. membrane blebbing. In Ca^2+^ experiments, control neurons were first selected at 20X magnification if they displayed observable spontaneous Ca^2+^ transients and showed no signs of membrane blebbing.

##### High-speed 3D SIM

Experiments were performed at the Advanced Imaging Center of the Janelia Research Campus on a custom-built SIM microscope that enables rapid acquisition of optical sections by employing a liquid crystal spatial light modulator with a rapid-switching variable retarder to efficiently alter the grating pattern, as described previously (Kner et al., 2009, Li et al., 2015). Images were collected using a 100× 1.46 NA Zeiss objective using a 560 nm excitation laser, an exposure time of 20 sec, pixel size of 0.04 µm, voxel depth of 0.15 µm, and 0.1 µm step size at a frame interval of 15-20 sec, depending on the number of slices, ranging from 19 to 21. This interval included a 7-10 sec imaging period and an equal recovery period to minimize the rate of photo-bleaching and extend the imaging duration to quantitate long-lived spinules. Notably, the enhanced acquisition speed compared to standard SIM results in fewer motion artifacts. Raw, rapid SIM datasets were reconstructed using a custom algorithm provided by Janelia researchers, and reconstructions were then imported into ImageJ-Fiji to convert from 32- to 16-bit images for further analysis.

##### ‘Enhanced resolution’ confocal microscopy

In instances that required endogenous protein labeling, less photo-bleaching, and/or increased speed than could be achieved by SIM, we utilized enhanced resolution confocal microscopy. Images were acquired using a Nikon A1R with resonant scanning and a GaAsP multi-detector unit using a 1.45 NA Plan Apo Lambda 100X oil objective and 488 and 561 nm excitation lasers. To approach the system’s optical limit, we adjusted the pinhole to 0.6 airy units, selected a pixel size of 0.06 µm (1/2 nyquist), and over-sampled in z by using step-size from 0.1 to 0.12 µm. We finally performed post-acquisition 3D deconvolution using Nikon Elements software 5.01, in automatic mode, to further enhance the resolution. For the experiments performed on neurons expressing mRuby and PSD95-specific intrabodies, a total of 41 to 43 slices were acquired with averaging at a frame interval between 8.2 and 8.47 sec. Bleach correction was applied to both channels using ImageJ-Fiji. For experiments performed on neurons expressing mRuby and GCAMP6s, 25 slices were acquired with averaging at a frame interval of 2.2 sec. In these datasets, only the red channel was deconvolved and bleach corrected, and the raw GFP MFI was utilized in the Ca^2+^ ΔF/F_0_ quantitation.

#### Image analysis and statistical analysis

For each analysis, cells were derived from two independent dissociated mouse neuron culture preparations. Large volumetric SIM datasets were obtained from two neurons, and smaller volumetric datasets obtained by enhanced resolution microscopy were from 8-11 neurons, i.e. 8 neurons for PSD imaging, 11 vehicle-treated and 10 BAPTA-AM-treated neurons for Ca^2+^ imaging. The numbers of dendritic spines and spinules analyzed in each condition are indicated in figure captions. Spines were excluded from further analysis if their entirety was not captured within the field of view throughout the imaging duration. Quantification of spine class, spine and spinule volumes, as well as number and lifespan of spinules over time was performed in 3D using IMARIS imaging analysis software versions8.4.1-9.1.2 (Bitplane). Three-dimensional volume view videos were also constructed using IMARIS. Spines closely associated with adjacent labeled neuronal elements were excluded from volume analysis by necessity. A mushroom spine was defined as one with a large head and thin neck, while a filopodia had a long, thin dendritic projection with no head, and thin spines displayed a long thin neck and small head (Hering and Sheng, 2001). A transitioning spine was defined as one that changed class, e.g. from filopodia to thin, during the imaging duration. Since mushroom spines predominantly formed spinules, we excluded other spine subtypes from subsequent in-depth analyses.

A spinule was generally defined as a thin projection of the mushroom spine membrane that was at least 0.225 µm in length. An individual spinule was defined as one that existed in a single frame or in consecutive frames, and the lifespan was defined as the number of frames in which a spinule was detected at a specific location on the spine head multiplied by sec per frame. Spinules lengths were measured from base to tip and widths measured at the narrowest point of the spinule from frame to frame. We observed that the widths of spinule-like projections were often not uniform along their length and also undulated over time, so we chose to include thicker structures (< 0.3 µm), provided the length was greater than width. Mean length, width, and volume measurements represent the average across all frames in which a spinule appeared consecutively. Individual tracking of spinule lifespans over time was achieved in 3D using IMARIS and spinule length measurements obtained using IMARIS Measurement Pro module or Image J-Fiji, while width measurements were obtained using Image J-Fiji.

##### Spinule recurrence

We first normalized the spinule origination point across the mushroom spine head to objectively quantitate this recurrence, considering that spines can rapidly change their shape and size and exhibit dynamic motility, including changes in spine head location relative to the dendritic shaft. In projection images, we traced the outline of the spine head ~0.1 µm from the edge at each time-point and generated single frame kymographs (plot profiles) of fluorescence intensity, with a peak in intensity signifying a spinule intersection and origination point. The length of each plot profile was then normalized on a generic unit scale of 0-100 and used to generate spatio-temporal plots to track spinule recurrence (within 10 units) in non-continuous frames in each spine. One outlier spinule had a lifespan that equaled or exceeded the duration of imaging, so recurrence could not be assessed, and it was excluded from further analysis.

##### Spinule assessment relative to the PSD

In addition to spinule lengths, the distance from the ‘spinule origination’ or base of the spinule to the nearest edge of the PSD were measured over time in 3D using the IMARIS measurement pro tool. The PSD edge was defined by GFP fluorescence detected in neurons expressing PSD95-specific GFP-tagged intrabodies, and mRuby fluorescence non-specifically labeled neuronal elements. To follow PSD volume and mean number of PSD fragments in individual dendritic spines, we utilized IMARIS surfaces with tracking and based on absolute GFP intensity on bleach-corrected images. The distinguishing features of simple versus segmented PSDs have been described previously *(18)*, and we additionally had to assess their changing features over time. Hence, PSDs were classified as partitioned if they displayed a moderate to high degree of segmentation and/or fragments, i.e. in ≥ 50% of frames. PSDs were classified as fragment-positive if distinctive fragments were detected ≥ 10% of frames. The mean and maximum number of fragments per frame was calculated objectively using IMARIS surfaces with tracking using a consistent threshold based on absolute GFP intensity.

##### Calcium transients

For experiments performed on neurons expressing both the mRuby cell fill and GCAMP6s, only the red channel was deconvolved, a maximum intensity projection generated in the z plane, and alignment of the red channel performed using NIS Elements. Regions of interest (ROIs) were drawn using the ROI editor within the dendritic shaft adjacent to the spine, spine head, and spinules, where possible. Those neuronal elements in close contact with other neuronal structures, which could not be adequately differentiated, were excluded from further analysis. ROIs were adjusted frame by frame and mean GFP MFI (raw AU) was calculated in NIS Elements for each of the 137 frames. From the MFI values, the ratio of ΔF/F_0_ was calculated by first averaging the values from four to five frames collected during an early period of inactivity to calculate F_0_. ΔF = F(t) - F_0_. Ca^2+^ peaks were defined as ΔF/F_0_ >0.50 or a 50% fluorescence increase over baseline. An individual peak ΔF/F_0_ maxima was defined as the highest value in a string of values above 0.5, and the peak duration was defined as the number of seconds spanned by a string of ascending and then descending values (over 0.50) around the maxima. Mean and maximum Ca^2+^ peak ΔF/F_0_ per spine refers to the average or maxima of all peaks detected in that spine over the 300 s imaging duration.

##### Statistics

Statistical analysis was performed using GraphPad Prism Version 7.0 (Graphpad Software). To determine the appropriate statistical text, normality of distribution was first tested using the D’Agostino-Pearson test. Parameters, such as spinule number and cumulative lifespan, between two independent groups were compared using two-tailed unpaired student t tests for normally distributed or Mann-Whitney test for non-normally distributed parameters between spines, e.g. the mean and maximum number of PSD fragments per frame. A two-tailed paired Student’s t test was applied to normally distributed data assessed within spines, e.g. the number of short-versus long-lived spinules per spine. For comparisons involving more than two independent groups, One-way ANOVA was applied to normally distributed data or Kruskal Wallis and Dunn’s Multiple Comparison test for non-normally distributed data to compare each group. To determine correlation between two variables, normally distributed data was assessed using the Pearson’s test for correlation, while the Spearman’s test was utilized for non-normally distributed data (both two-tailed). The Fisher’s Exact test was used to analyze contingency tables of two subgroups, long-lived spinule positive and negative spines, and two outcomes, e.g. non-partitioned *versus* partitioned PSDs or non-fragmented *versus* fragmented PSDs. The number of cells and spines and/or spinules (n) is reported in the figure legends. Data are represented as mean ± SEM. *P<0.05, **P<0.01, ***P<0.001, ****P<0.0001.

### Supplementary video legends

Video 1. Spinule recurrence on mushroom spines

Rapid SIM and 3D reconstructions depict spinules recurring at the same topographical location on a mushroom spine head and repeatedly extending and retracting along an axon over the 1000 sec duration of imaging.

Video 2. Typical spinule lifespans and morphologies

Rapid SIM of a large mushroom spine displaying multiple short-lived spinules with vermiform and tapered morphologies as well as long-lived spinules of varying shapes over 1000 sec.

Video 3. Differential short-lived and long-lived spinule dynamics

Wireframe 3D model from rapid SIM of a mushroom spine exhibiting one long-lived, pinch-waisted, stable spinule (lifespan≥1000 sec) and one short-lived, dynamic spinule (lifespan=40 sec).

Video 4. PSD fragment trafficking within a long-lived spinule

Enhanced resolution confocal imaging of a mushroom spine displaying a complex, partitioned PSD and a stable, long-lived spinule. A PSD fragment (green) travels to the base and into the spinule.

Video 5. Long-lived spinules are associated with complex PSDs

Enhanced resolution confocal imaging of a mushroom spine exhibiting a long-lived spinule that originates from a gap in the highly complex, fragmented, dynamic PSD.

Video 6. PSD fragment trafficking in a filopodia-like spinule

A PSD fragment (green) localizes to the tip of a rarely occurring, long, filopodia-like spinule, that originates relatively remote from a complex PSD. As the spinule retracts, the fragment travels along its length toward the spine head.

Video 7. Ca^2+^ puncta appearing in spinules during spine head transients

Enhanced resolution imaging of a GCAMP6s-expressing control neuron shows a spine with frequent, high amplitude Ca^2+^ peaks, multiple spinules, and Ca^2+^ puncta localizing to a long-lived spinule during spine head transients.

Video 8. Spines with frequent high-amplitude Ca^2+^ transients and many spinules

Representative control spines display frequent, high amplitude Ca^2+^ peaks and many spinules. Ca^2+^ puncta within a long-lived spinule is synchronized with Ca^2+^ transients in the spine head and dendritic shaft.

Video 9. Spinule number decreased following Ca^2+^ depletion

Enhanced resolution confocal imaging of representative mushroom spines from a GCAMP6-GFP-expressing BAPTA-AM-treated neuron that display dampened Ca^2+^ transients and very few spinules, compared to controls.

